## Supplementary information for "G protein-coupled receptor 17 inhibits glucagon-like peptide-1 secretion via a Gi/o-dependent mechanism in enteroendocrine cells"

### Supplementary Figure 1

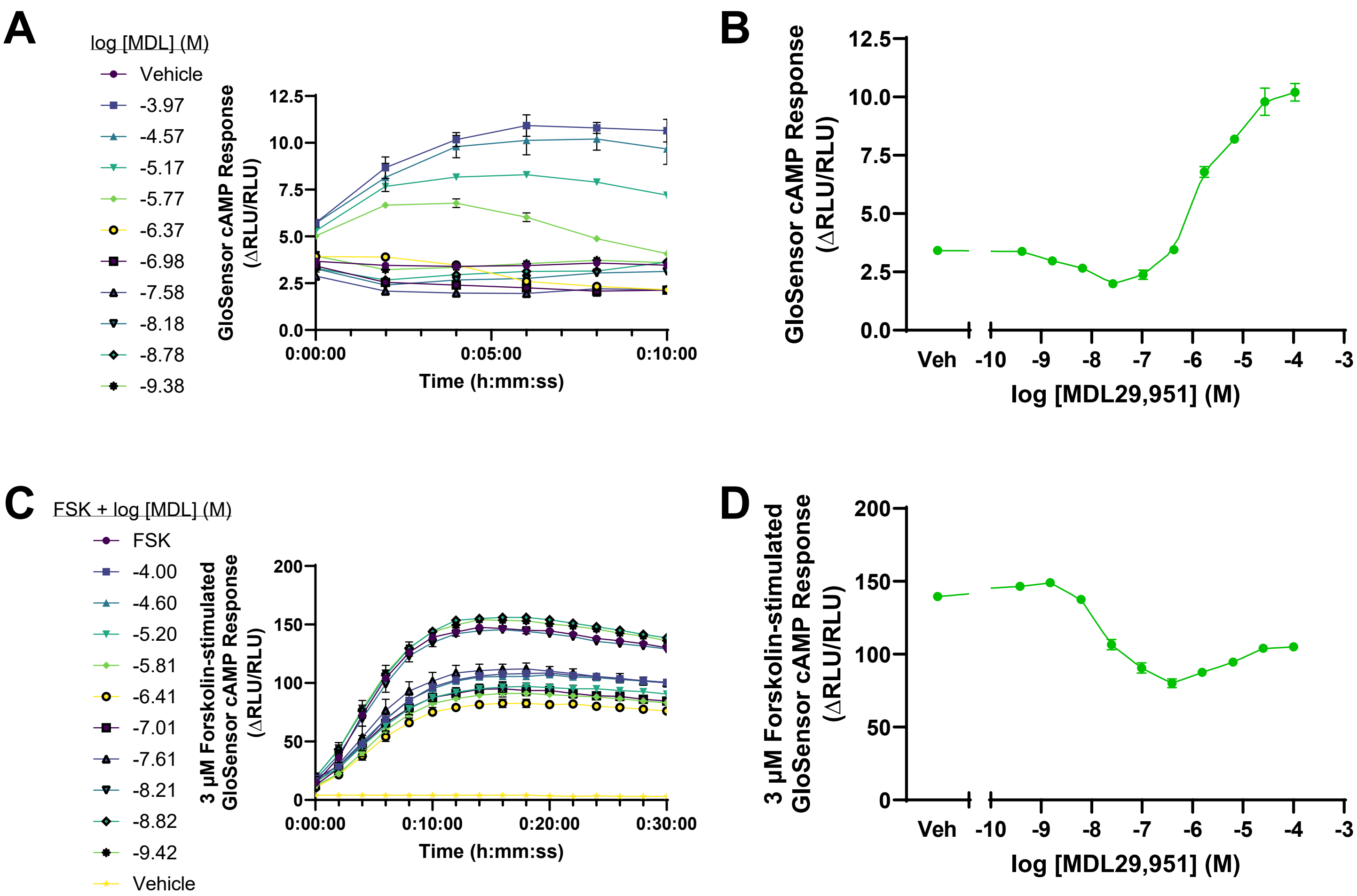


**Figure S1. Human GPR17 long isoform modulates cAMP in GLUTag cells. (A)** GloSensor cAMP luminescence responses were measured in GLUTag cells transiently transfected with hGPR17L and pGloSensor-cAMP-22F. Cells were treated with 500 μM IBMX and then stimulated with indicated concentrations of MDL29,951 and the luminescence response was measured every two minutes for ten minutes. **(B)** Concentration-response curve for MDL29,951 effects on GloSensor cAMP response four minutes after treatment. **(C)** GloSensor cAMP luminescence responses were subsequently measured in the same cells after stimulation with 3 μM forskolin. **(D)** Concentration-response curve for MDL29,951 effects on GloSensor cAMP signal following treatment with 3 μM forskolin. Data points represent the average responses for measurements between 16 and 30 minutes after forskolin stimulation. Data represent mean±SEM of two independent experiments performed in duplicate.

### Supplementary Figure 2

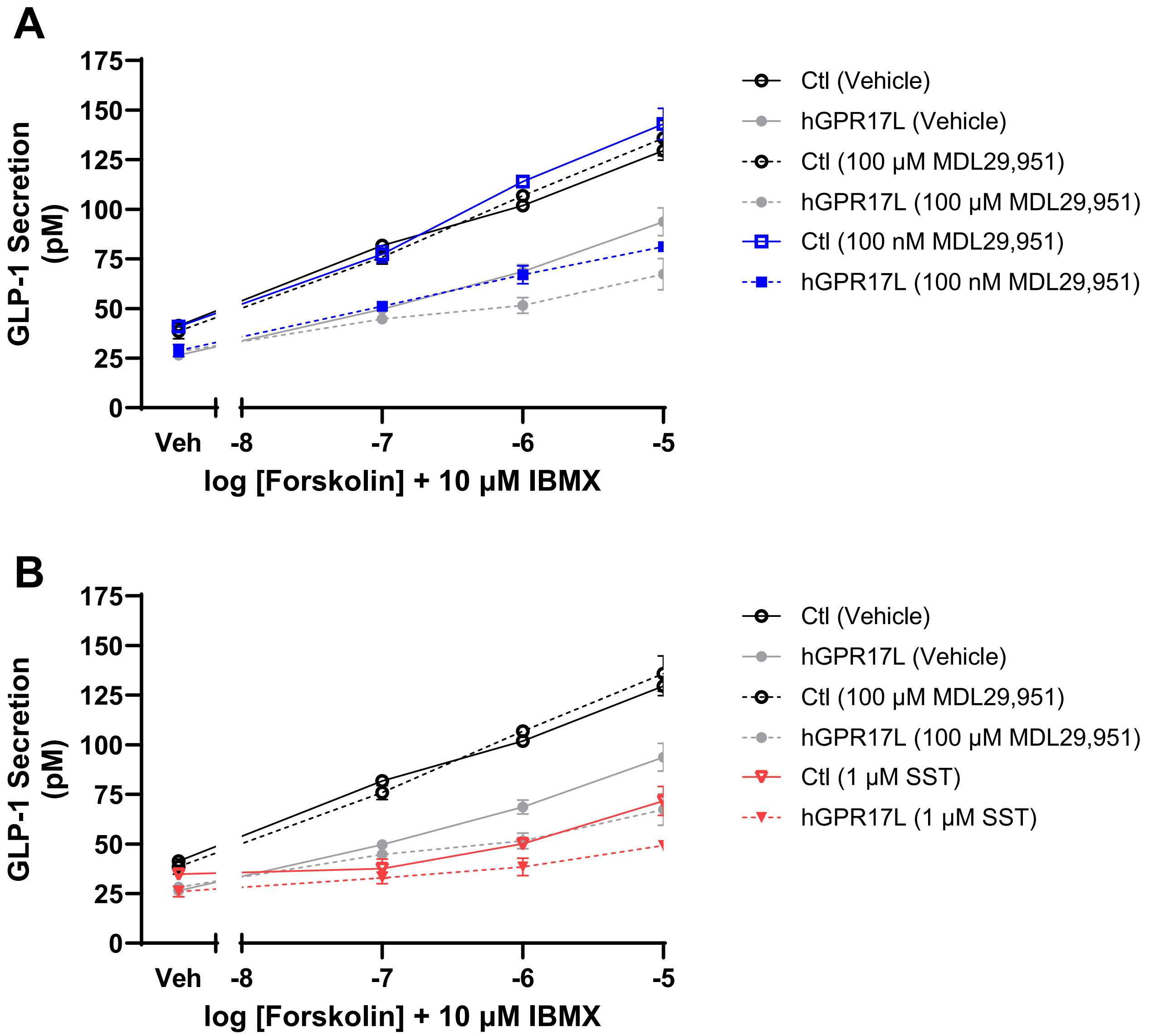


**Figure S2. Somatostatin negatively regulates GLP-1 secretion from GLUTag cells.** GLP-1 secretion was measured from control or hGPR17L-expressing GLUTag cells that were treated with **(A)** 100 nM MDL29,951 or **(B)** 1 µM somatostatin (SST) together with varying concentrations of forskolin and 10 µM IBMX. GLP-1 secretion in response to vehicle or 100 µM MDL29,951 treatments from Figure 3C are shown for reference comparison. Data are mean ± SEM of two independent experiments.

### Supplementary Figure 3 and Table S1

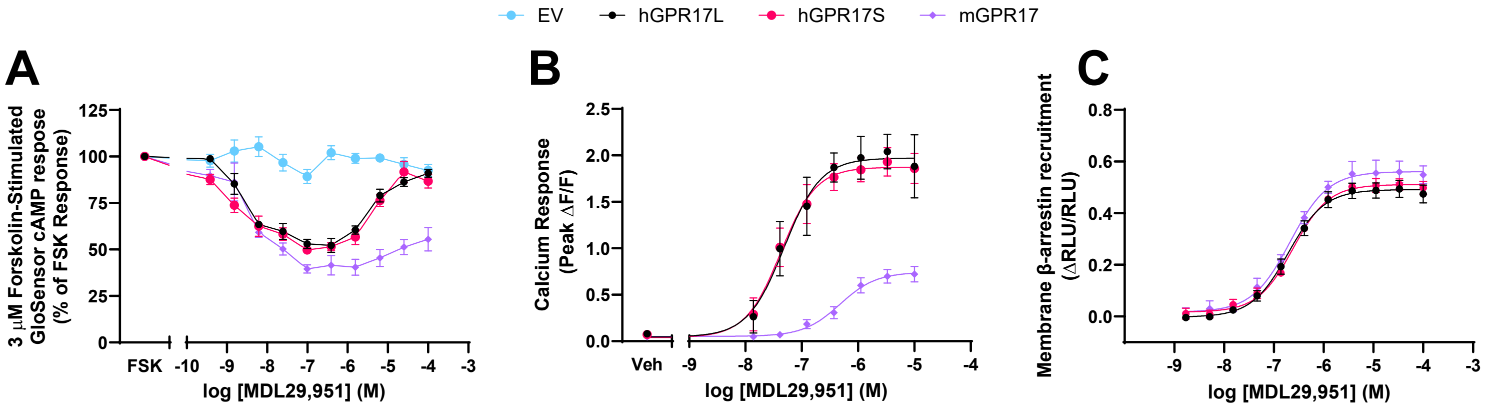


**Figure S3. Human GPR17 and mGPR17 have distinct second messenger signaling profiles in HEK293 cells.** HEK293 cells were transiently transfected with hGPR17L, hGPR17S, mGPR17, or empty vector (EV) and second messenger signaling regulation was measured. Concentration-response curves for MDL29,951 effects on **(A)** 3 μM forskolin-stimulated GloSensor cAMP signal, **(B)** calcium mobilization, and **(C)** membrane β-arrestin recruitment. Data represent mean±SEM of three to four independent experiments.

**Table S1. Human GPR17 and mGPR17 have distinct second messenger signaling profiles in HEK293 cells.** Data represent summaries of the potency and efficacy measures for concentration-response analyses of MDL29,951 effects on 3 μM forskolin-stimulated GloSensor cAMP signal, calcium mobilization, and membrane β-arrestin recruitment that were reported in Figure S3A, S3B, and S3C, respectively. Data represent mean±SEM of three to four independent experiments. Efficacy measures were statistically analyzed by one-way ANOVA with Dunnett's multiple comparisons test as compared to the hGPR17L transfection condition. * $p < 0.05$, ** $p < 0.01$, *** $p < 0.001$. EV, empty vector. nd, not determined.

| | cAMP | | | | Calcium | | β-arrestin | |
|---|---|---|---|---|---|---|---|---|
| | Potency | Maximum Inhibition | Potency | Effect of 100 μM MDL | Potency | Emax | Potency | Emax |
| Receptor | $IC_{50}$ (nM) (pIC50) | (% Inhibition) | $EC_{50}$ (μM) (pEC50) | (% Inhibition) | $EC_{50}$ (nM) (pEC50) | (Mean Peak ΔF/F) | $EC_{50}$ (nM) (pEC50) | (Peak ΔRLU/RLU) |
| hGPR17L | 3.0 (8.52±0.08) | 48±2.9 | 4.0 (5.40±0.09) | 9.0±2.1 | 51 (7.29±0.11) | 1.92±0.22 | 190 (6.72±0.09) | 0.50±0.03 |
| hGPR17S | 1.5 (8.83±0.09) | 48±2.2 | 3.6 (5.45±0.07) | 13±3.7 | 43 (7.37±0.10) | 1.84±0.11 | 240 (6.62±0.08) | 0.50±0.04 |
| mGPR17 | 3.3 (8.48±0.20) | 61±3.4* | 18 (4.74±0.23) | 45±6.3*** | 490 (6.31±0.05) | 0.70±0.09** | 200 (6.69±0.10) | 0.54±0.03 |
| EV | nd | nd | nd | 7.5±3.2 | nd | nd | nd | nd |

### Supplementary Figure 4

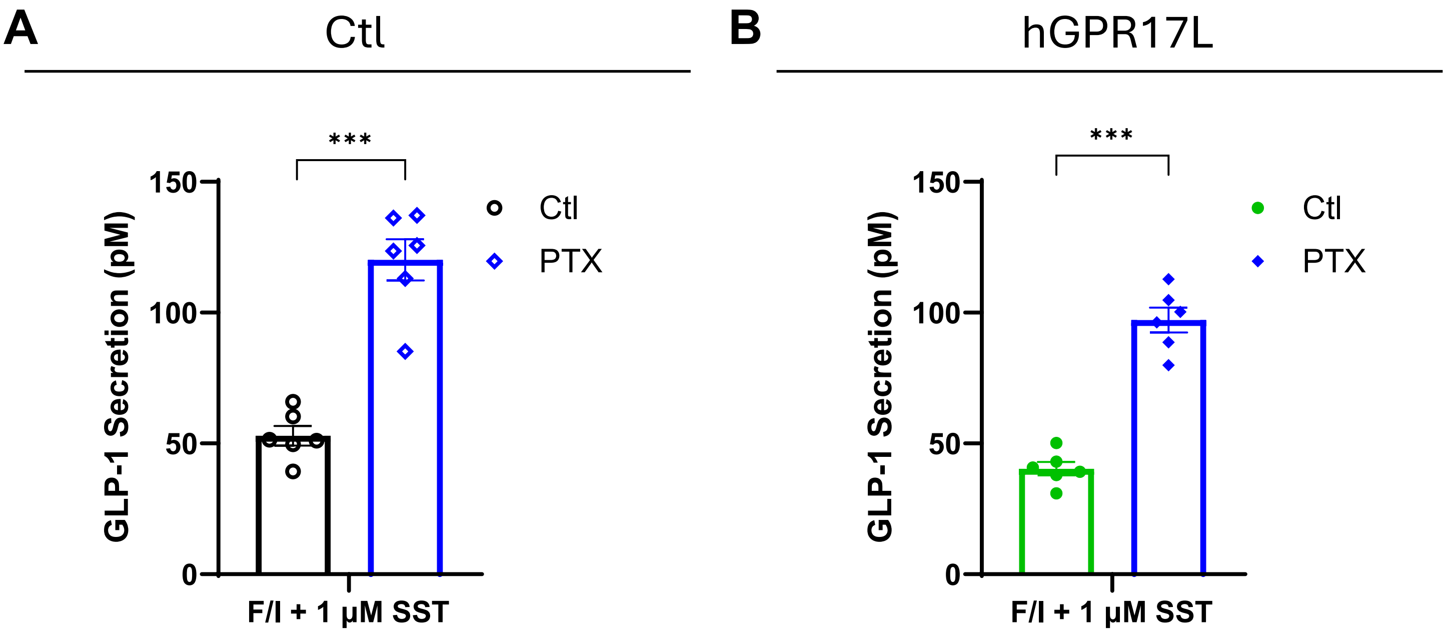


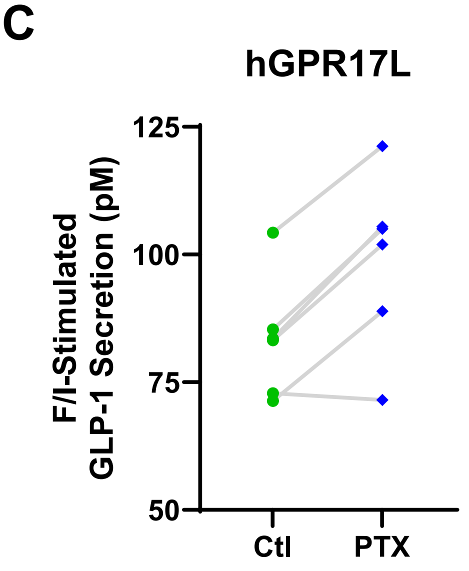


| Transduction | hGPR17L | hGPR17L | |
|---|---|---|---|
| Treatment 1 | Ctl | PTX | |
| Treatment 2 | Vehicle | Vehicle | |
| Treatment 3 | 10 μM F/I | 10 μM F/I | |
| | GLP-1 Secretion (pM) | | Δ |
| Rep1 | 104.3 | 121.2 | 17.0 |
| Rep2 | 83.2 | 102.0 | 18.8 |
| Rep3 | 72.9 | 71.6 | -1.3 |
| Rep4 | 83.5 | 105.5 | 22.0 |
| Rep5 | 71.3 | 88.9 | 17.6 |
| Rep6 | 85.3 | 105.0 | 19.7 |
| Mean | **83.4** | **99.0** | **15.6** |
| SEM | 4.8 | 6.9 | 3.5 |

**Figure S4. Pertussis toxin attenuates somatostatin-mediated inhibition of GLP-1 secretion and hGPR17L-mediated constitutive inhibition of GLP-1 secretion.** GLUTag cells were transduced with **(A)** control adenovirus or **(B)** adenovirus encoding hGPR17L, and GLP-1 secretion was measured in cells that were treated as indicated together with either control or 100 ng/mL PTX. Data represent mean ± SEM for five or six independent experiments and were analyzed with unpaired t tests comparing matched control and PTX treated conditions. ***, $p <$ 0.001. **(C)** Panel and table of data from Figure 5B demonstrating effects of PTX on F/I-stimulated GLP-1 secretion from matched experiments in GLUTag cells expressing hGPR17L.

### Supplementary Figure 5

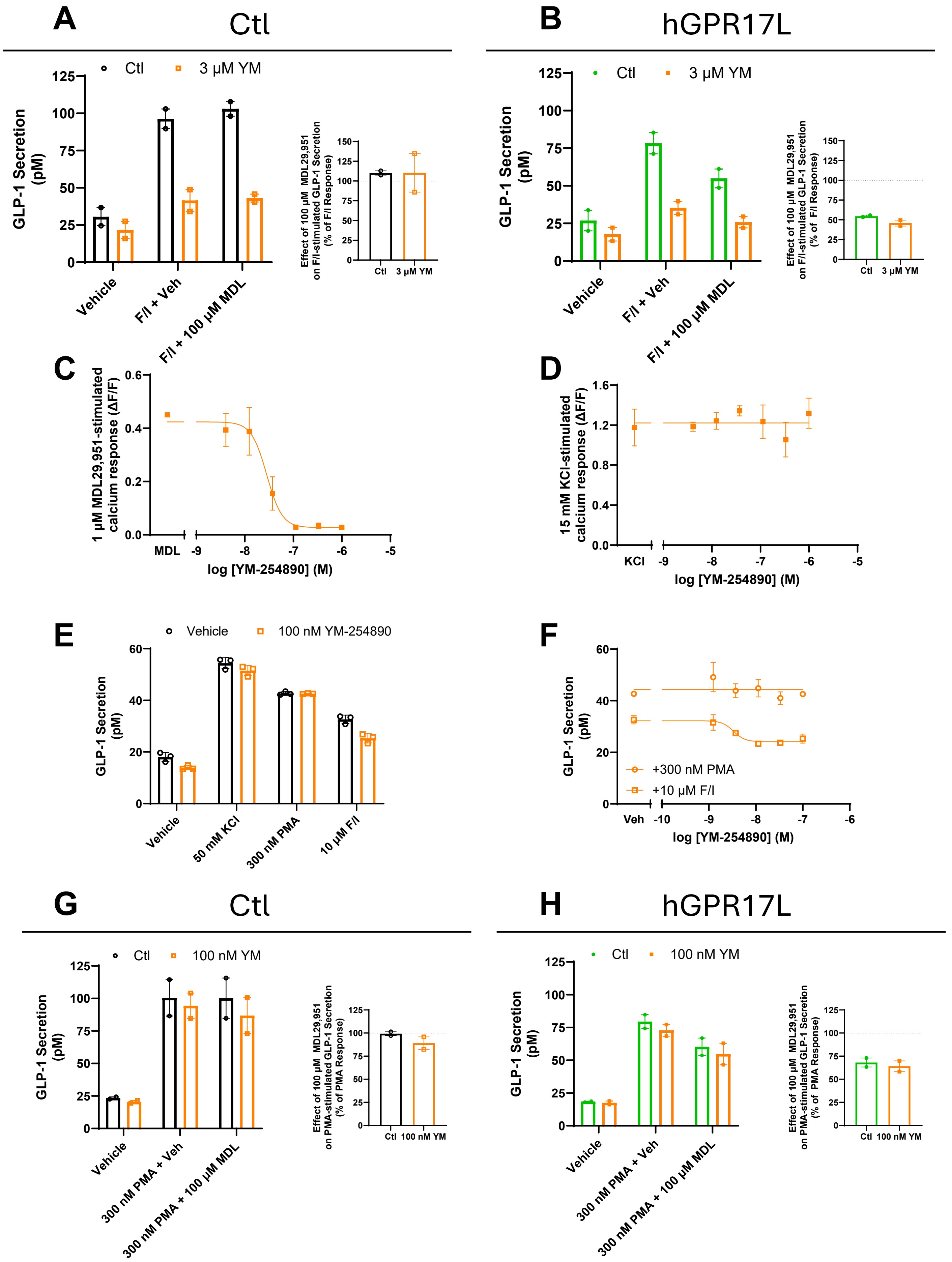

**Figure S5. YM-254890 selectively inhibits GPR17 agonist-mediated calcium flux but has no effect on hGPR17L-mediated inhibition of GLP-1 secretion.** GLUTag cells were transduced with **(A)** control adenovirus or **(B)** adenovirus encoding hGPR17L and GLP-1 secretion was measured in cells that were treated as indicated together with either control or 3 μM YM-254890. Data represent mean±SEM for two independent experiments performed in duplicate. GLUTag cells expressing hGPR17L were treated with indicated concentrations of YM-254890 and stimulated with **(C)** 1 μM MDL29,951 or **(D)** 15 mM KCl and calcium mobilization was measured. Data represent mean±SD of a single experiment performed in triplicate. GLP-1 secretion was measured from GLUTag cells that were stimulated as indicated in the presence of **(E)** control or 100 nM YM-254890 treatment or **(F)** indicated concentrations of YM-254890. Data represent mean±SD of a single experiment performed in triplicate. GLUTag cells were transduced with **(G)** control adenovirus or **(H)** adenovirus encoding hGPR17L and GLP-1 secretion was measured in cells that were treated as indicated together with either control or 100 nM YM-254890. Data represent mean±SEM for two independent experiments performed in duplicate.

### Supplementary Figure 6

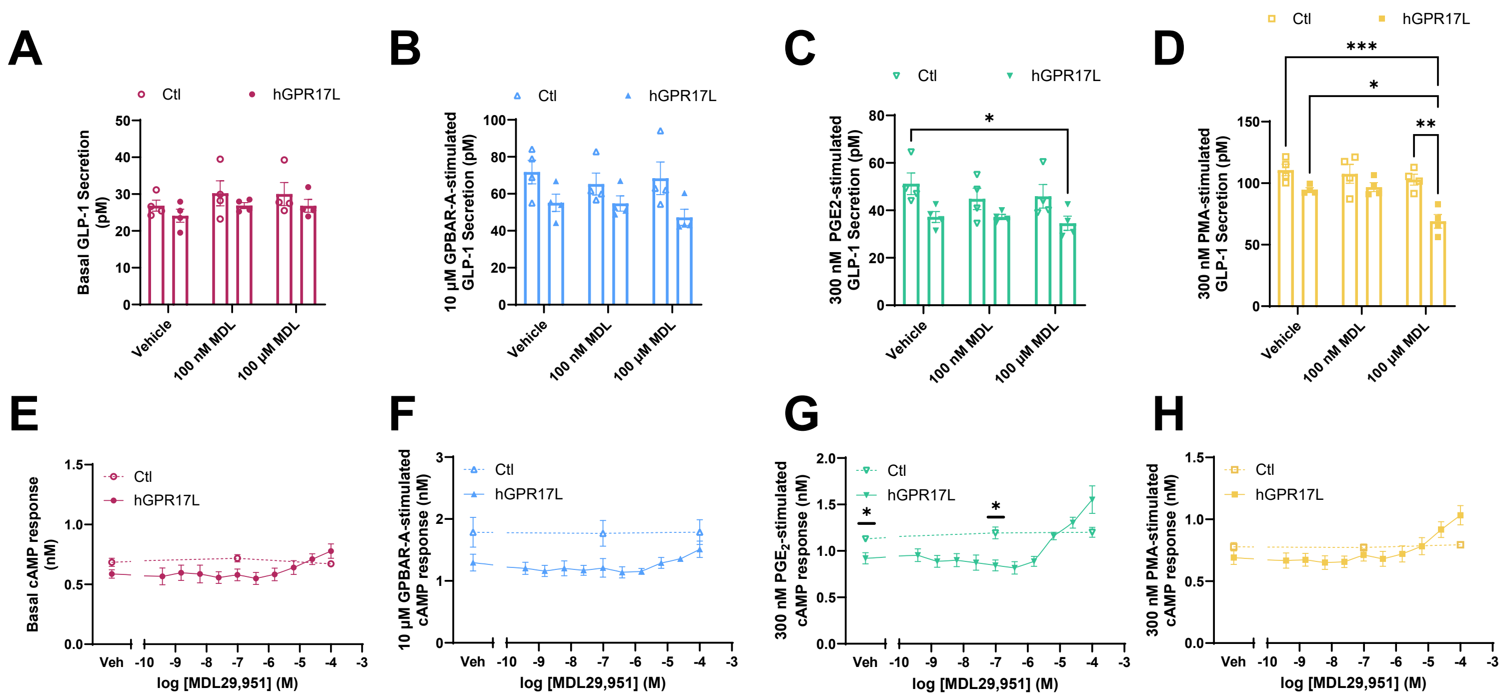


**Figure S6. Human GPR17 long isoform distinctly regulates GLP-1 secretion and cAMP signaling in a manner that is dependent on co-stimulation conditions in GLUTag cells.** GLUTag cells were transduced with control adenovirus or adenovirus encoding hGPR17L and GLP-1 secretion was measured in response to treatment with **(A)** vehicle, **(B)** 10 µM GPBAR-A, **(C)** 300 nM $PGE_2$, or **(D)** 300 nM PMA together with vehicle, 100 nM MDL29,951, or 100 µM MDL29,951. Data represent mean±SEM of four independent experiments performed in triplicate and were analyzed using one-way ANOVA with Sidak's post hoc test. *, $p < 0.05$, **, $p < 0.01$, ***, $p < 0.001$. Cyclic AMP was measured under **(E)** vehicle, **(F)** 10 µM GPBAR-A, **(G)** 300 nM PGE2, or **(H)** 300 nM PMA stimulation conditions in combination with the indicated concentrations of MDL29,951 in control or hGPR17L-expressing GLUTag cells. Data represent mean±SEM for two to four independent experiments performed in triplicate and were analyzed by unpaired t test comparing matching treatment conditions between control and hGPR17L-expressing cells. *, $p < 0.05$.
